# Accurate Identification of Active Transcriptional Regulatory Elements from Global Run-On and Sequencing Data

**DOI:** 10.1101/011353

**Authors:** Charles G. Danko, Stephanie L. Hyland, Leighton J. Core, Andre L. Martins, Colin T. Waters, Hyung Won Lee, Vivian G. Cheung, W. Lee Kraus, John T. Lis, Adam Siepel

## Abstract

Identification of the genomic regions that regulate transcription remains an important open problem. We have recently shown that global run-on and sequencing (GRO-seq) with enrichment for 5'-capped RNAs reveals patterns of divergent transcription that accurately mark active transcriptional regulatory elements (TREs), including enhancers and promoters. Here, we demonstrate that active TREs can be identified with comparable accuracy by applying sensitive machine-learning methods to standard GRO-seq and PRO-seq data, allowing TREs to be assayed together with transcription levels, elongation rates, and other transcriptional features, in a single experiment. Our method, called <u>d</u>iscriminative <u>R</u>egulatory <u>E</u>lement detection from <u>G</u>RO-seq (dREG), summarizes GRO-seq read counts at multiple scales and uses support vector regression to predict active TREs. The predicted TREs are strongly enriched for marks associated with functional elements, including H3K27ac, transcription factor binding sites, eQTLs, and GWAS-associated SNPs. Using dREG, we survey TREs in eight cell types and provide new insights into global patterns of TRE assembly and function.

---

Transcriptional regulatory elements (TREs), such as promoters, enhancers, and insulators, are critical components of the genetic regulatory programs of all organisms^1^. These elements regulate gene expression by facilitating or inhibiting chromatin decompaction, transcription initiation, and the release of RNA polymerase II into productive elongation, as well as by maintaining the three-dimensional architecture of the nucleus. TREs enable complex, cell-type-and condition-dependent patterns of gene expression that contribute to nearly all biological processes.

Since the completion of high-quality gene catalogs for humans and most model organisms, the comprehensive identification and characterization of TREs has emerged as a major challenge in genomic research. These elements are most effectively identified using high-throughput genomic assays that provide indirect evidence of regulatory function, such as chromatin immunoprecipitation of bound transcription factors (TFs), histone modifications, or DNase-I hypersensivity and sequencing (DNase-seq)^2–4^. However, the methods currently in wide use have important limitations. For example, ChIP-seq requires a high-affinity antibody for the targeted TF or histone modification of interest and must be performed separately for each target. Likewise, assays that measure chromatin accessibility or histone modifications provide only circumstantial evidence that the identified sequences are actively participating in transcriptional regulation^5^. Even STARR-seq, a clever high-throughput reporter gene assay, identifies a subset of regions that are inactive *in situ*, because the reporter assay is independent of local chromatin structure and normal genomic context^6^.

Recently it has become clear that a defining characteristic of active TREs is that they are associated with local transcription. Enhancer-templated non-coding RNAs, or eRNAs, previously thought to be rare^7,8^, have recently been associated with thousands of stimulus-dependent enhancers^9^. Like active promoters, these enhancers exhibit transcription initiation in opposing directions on each strand, a phenomenon called divergent transcription^10–12^. This characteristic pattern can be a powerful tool for the identification of active TREs in a cell-type specific manner^9,13–15^. Methods that measure the production of nascent RNAs, such as Global Run-On and sequencing (GRO-seq)^10^ and its successor, Precision Run-On and sequencing (PRO-seq)^16^, are particularly sensitive for detecting these transient enhancer-associated RNAs, because they detect primary transcription before unstable RNAs are degraded by the exosome^14,17^. Recently, we have shown that an extension of GRO-seq that enriches the nuclear run-on RNA pool for 5'-7meGTP-capped RNAs, called GRO-cap, further improves sensitivity for eRNAs, and can be used to identify tens of thousands of transcribed enhancers and promoters across the genome^18,19^.

Here we introduce a new computational method for identifying transcribed TREs directly from standard GRO-seq or PRO-seq data, with nearly the accuracy of the custom GRO-cap procedure. Our method, called discriminative Regulatory Element detection from GRO-seq (dREG), makes use of a novel, multiscale summary of GRO-seq or PRO-seq read counts, and then employs support vector regression^20^ (SVR) to detect the characteristic patterns of transcription at TREs. dREG was trained using data generated by GRO-cap^19^, which defines a high-confidence set of TREs associated with divergent transcription. The resulting dREG method allows high-quality predictions of TREs for any cell type with existing GRO-seq or PRO-seq data. Here, we apply the method to four cell types for which data was previously available and four for which we provide new data. Combining these predictions with ones based on chromatin modifications, CTCF ChIP-seq, and DNase-seq, we find that the predicted TREs fall into four major classes. One class, which is defined by a strong dREG signal, is also enriched for H3K27 acetylation (H3K27ac), binding sites for sequence-specific TFs, eQTLs, and GWAS-associated SNPs. Together these findings support a hierarchical model for transcriptional regulation, with transcribed TREs identified using dREG most directly reflecting cell-type-specific activation. Genome browser tracks describing our predictions and the dREG software package are freely available.

## Results

### Design, Training, and Validation of dREG

TREs identified by chromatin modifications and DNase-seq exhibit clear signs of bidirectional transcription, which are apparent from PRO-seq data (Fig. 1). We devised a machine-learning approach, called dREG, to identify TREs, including both promoters and distal enhancers, from standard GRO-seq or PRO-seq data. The key to our method is a feature vector that summarizes the patterns of aligned GRO-seq reads near each candidate element at multiple scales. This feature vector consists of read counts for windows ranging in size from 10 bp to 5 kbp, standardized using the logistic function (Fig. 2A). The feature vector is passed to a standard SVR, which scores sites with high PRO-seq signal for similarity to a training set of TREs (see methods for details). To train our classifier, we used TREs identified from GRO-cap data^19^ as positive examples and regions of matched PRO-seq signal intensity lacking additional marks associated with TREs as negative examples. Various tuning parameters, including the number of windows, the window sizes, and the standardization strategy, were optimized by cross-validation (Supplementary Tables 1-2). After training and optimization, the program displayed excellent performance when applied to PRO-seq data for K562 cells (AUC= 0.98; Fig. 2B).

**Fig. 1:**
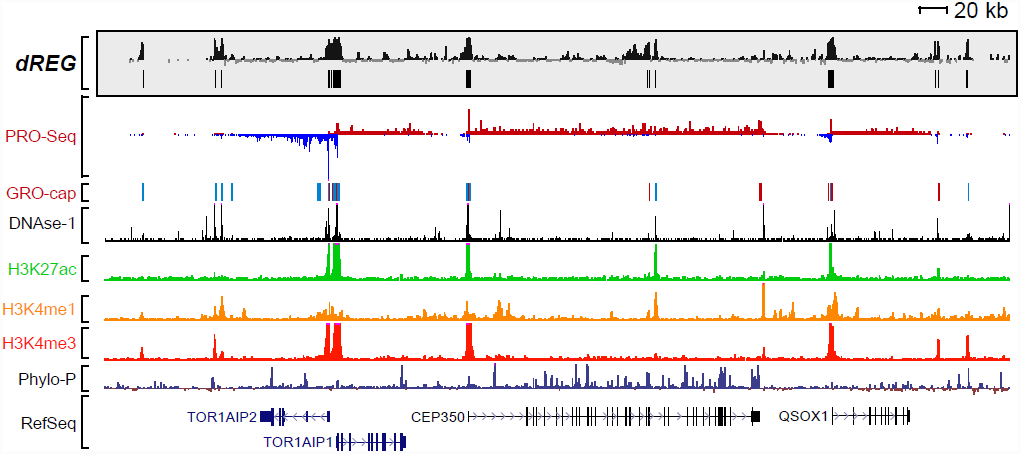
Browser shot demonstrating the dREG technique. Browser shot depicts raw dREG scores and dREG ‘peaks’ alongside PRO-seq, GRO-cap, DNase-I, and ENCODE ChIP-seq data for H3K27ac, H3K4me1, and H3K4me3.

**Fig. 2:**
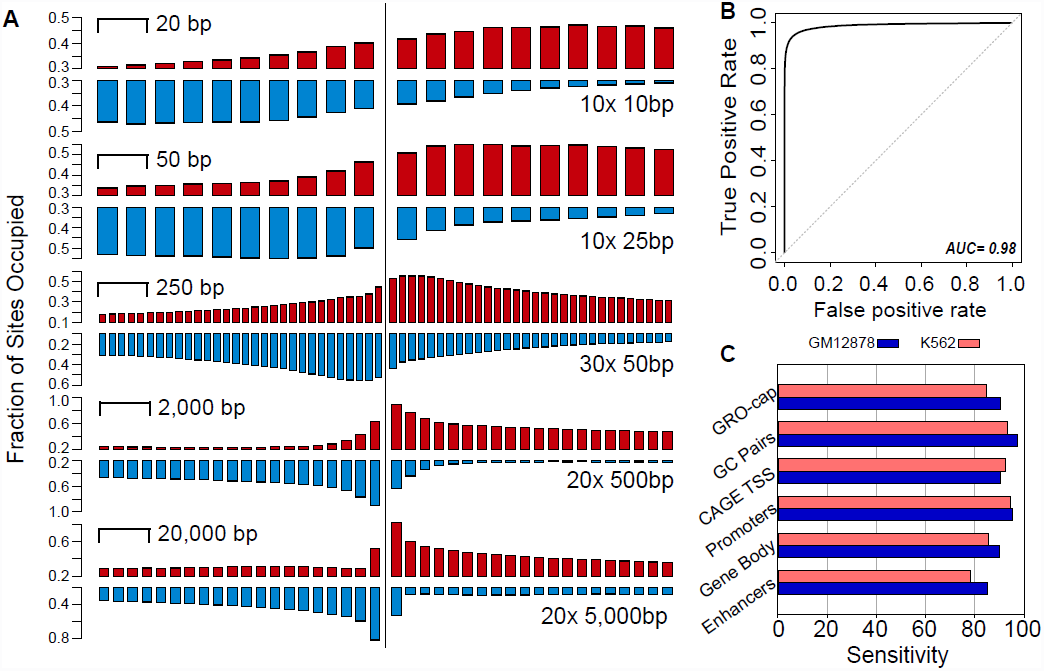
dREG model signal and validation. (A) The signal for +dREG TREs depicted as barcharts at decreasing window size (from top to bottom). Boxes represent consecutive, non-overlapping window sizes. The number of windows in the optimal feature vector, and the size represented by each bar, is shown at right. (B) ROC plot shows the accuracy of dREG at distinguishing regulatory TREs in a matched task (AUC= 0.98). (C) Genome-wide sensitivity for the indicated classes of regulatory elements at a 10% false discovery rate in K562 (pink) and GM12878 (blue). Enhancers and promoters reflect labels assigned by ChromHMM, and gene bodies were identified by intersecting GRO-cap peaks with RefSeq annotations.

We ran dREG to predict the location of transcribed regulatory regions genome-wide in K562 cells, adopting a prediction threshold that limits the genome-wide false discovery rate to 10%. At this threshold, we recovered 93% of 15,276 GRO-cap 'paired' sites (i.e., sites for which divergent Pol II initiation was detected in both directions), and 92% of 10,024 active transcription start sites detected by CAGE (Fig. 2C). Furthermore, we observed high sensitivity within the subsets of GRO-cap peaks overlapping promoters (94%), enhancers (78%), or gene bodies (85%) (Fig. 1,2C). In addition, we applied dREG to an independent cell type, GM12878 lymphoblastoid cells, without retraining the classifier. Based on GRO-cap data available for GM12878, dREG achieved similar performance in this cell type for all classes of regulatory elements tested (Fig. 2C), indicating that dREG generalizes well across cell types. Finally, we examined the sensitivity of dREG to sequencing depth and data quality and found that sensitivity is satisfactory with as few as 40M mapped reads and saturation is achieved by ~100M mapped reads (Supplementary Fig. 1). Together, these findings demonstrate that dREG accurately identifies active TREs across a broad spectrum of GRO-seq and PRO-seq data sets.

### Prediction of Transcriptional Regulatory Elements across Eight Cell Types

dREG enabled us to generate predictions of TREs for several cell types for which GRO-seq or PRO-seq data is available. We analyzed existing GRO-seq data sets for MCF-7, IMR90, GM12878, and AC16 cell lines^10,13,21–23^, as well as new data that we generated in four cell types analyzed by the ENCODE and Epigenome Roadmap projects, including K562, primary CD4+ T-cells, Jurkat leukemia cells, and HeLa carcinoma cells. For each of these new cell types, GRO-seq or PRO-seq libraries were produced, validated for library complexity, and sequenced to a depth of 53–375 million reads (Supplementary Table 3). The dREG model trained on K562 cells was applied to each data set. The dREG predictions for each cell type include ~30,000 TREs (21,232-37,947), covering ~1% of the human genome. Approximately half of these elements mark active promoters and half mark a subset of distal enhancers (Supplementary Fig. 2). The union of these predictions across all eight cell types includes 105,356 TREs, covering 3.7% of the human genome.

### Four Major Classes of Transcriptional Regulatory Elements

We enlarged our set of candidate regulatory elements by combining the TREs identified by dREG with putative TREs identified by two complementary methods: ChromHMM predictions of promoters, enhancers, and insulators^24^, and DNase-I hypersensitive sites (DHSs), which correlate broadly with TREs^25,26^. The ChromHMM predictions are based on genome-wide measurements of post-translational histone modifications (including, but not limited to H3K27ac, H3K4me1, H3K4me3, and H4K20me1) and CTCF binding, whereas the DHSs identify regions of ‘open’ chromatin where the DNA is accessible to DNase-I cleavage. For the DHSs, we used high-confidence DNase-I accessible sites, defined as the intersection of Duke and UW DHS predictions (Supplementary Fig. 3). After taking the union of these three sets of putative TREs (merging overlapping elements; see Methods), we labeled each TRE by the collection of methods that identified it (dREG, ChromHMM, and/or DNase-seq). This analysis focused on four cell types (K562, GM12878, HeLa, and CD4+ T-cells) for which data was available from all assays.

**Fig. 3:**
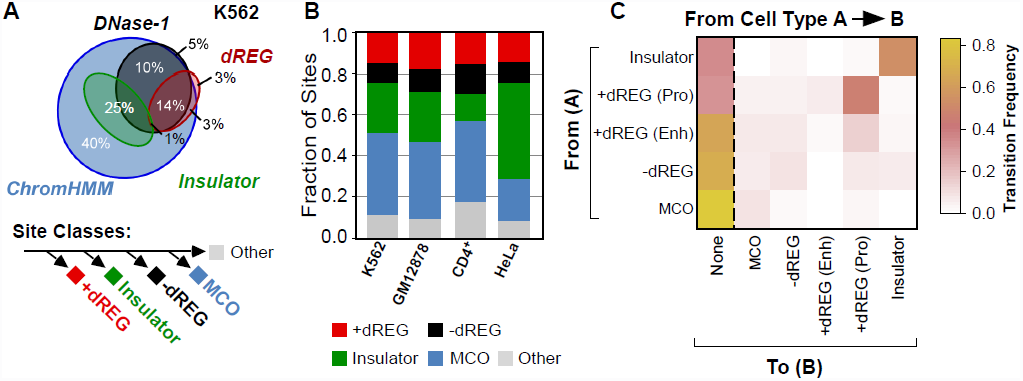
Comparison of putative TREs detected using dREG, DNase-I, and ChromHMM. (A) Four-way Venn diagram depicting the overlap between four classes of regulatory element. TREs were classified as +dREG, insulator, -dREG, or as modified chromatin only (MCO). Numbers give the rounded overall fraction of TREs that fall into the specified intersection. (B) The fraction of TREs in each of the four classes compared across four cell types (K562, GM12878, CD4+ T-cells, and HeLa carcinoma cells). (C) Heatmap denotes the median frequency of differences between cell types of the indicated classes of functional element.

Our analysis of the labeled TREs indicated that these three methods identified nested sets of elements, with the ChromHMM predictions being most inclusive, the DHSs largely forming a subset of the ChromHMM predictions, and dREG, in turn, generally narrowing those identified by DNase-seq to a smaller subset (Fig. 3A,B). Interestingly, the ChromHMM predictions of insulators showed limited overlap with DNase-seq or dREG predictions. Thus, we observe four main classes of TREs based on these methods: (1) actively transcribed TREs identified by dREG, DNase-seq, and ChromHMM, comprising 14–17% (depending on the cell type) of the merged set (+dREG); (2) ‘open’ but untranscribed TREs identified by DNase-seq and ChromHMM (excluding the insulator predictions) but not dREG, accounting for 10% (-dREG); (3) elements with histone modifications indicative of enhancers but that are untranscribed and display either weak (16.8%) or no (23.2%) evidence of DNase-I accessibility, accounting for 40% (Marked Chromatin Only; or MCO); and (4) the ChromHMM insulator predictions, which comprise 25% of all TREs (Insulator). Other combinations of assays account for only 1–5% of TREs, and can likely be attributed to experimental biases, false positive and/or false negative predictions. Notably, a follow-up analysis suggested that the absence of dREG predictions in the-dREG set does not reflect any inadequacy of sensitivity or sequencing depth (Supplementary Fig. 4). These observations indicate that dREG identifies a smaller collection of transcribed TREs that might have functional properties that distinguish them from sites predicted using chromatin modifications or DNase-seq alone.

**Fig. 4:**
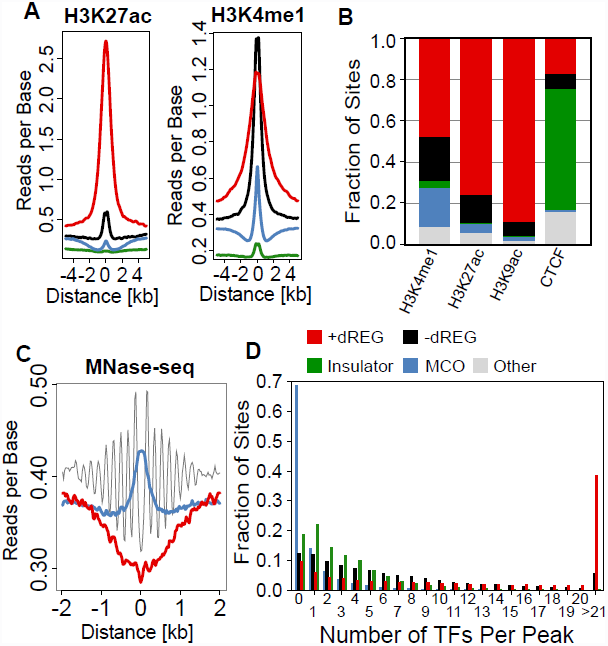
The enrichment of functional marks in each class of TRE. (A) Comparison of read-densities for H3K27ac (left) and H3K4me1 (right) in each class of functional element. (B) The fraction of ENCODE peak calls for the specified mark (H3K4me1, H3K27ac, H3K9ac, and CTCF) in each of the four classes. The ‘other’ category denotes peaks for the indicated mark falling outside of the putative TREs identified by other assays. (C) Distribution of MNase-seq reads in the +dREG (red), -dREG (black), and modified chromatin only classes (MCO; blue). (D) Histogram compares the number of transcription factors found in each of the four functional classes.

### Relationship between the Four TRE Classes across Cell Types

Although TREs fall into distinct classes within a cell type, we hypothesized that they might transition from one class to another across different cell types. As previous work suggests different rates of cell-type-specific expression at promoters and enhancers, we additionally separated +dREG into non-overlapping sets of promoters and enhancers using the ChromHMM labels, leading to a total of five classes per cell type. We then compared the frequencies of differences between TRE classes (and between sites that are ‘unmarked’, or not a TRE) across pairs of cell types for which all sources of data were publically available (K562, GM12878, CD4+ T-cells, and HeLa). For visualization, we generated a heatmap based on the median frequency with which differences in class assignment are observed across all cell pairs (Fig. 3C). This analysis revealed that the MCO-TREs are more frequently cell-type-specific compared with the other classes (78% transition from MCO to unmarked). By contrast, promoters and insulators are significantly more likely to be conserved across cell types than are other types of TREs. TREs that mark distal enhancers (including both +/-dREG classes) are commonly specific to a single cell type (63-64% transition to unmarked). The remaining 36-37% of distal enhancer TREs have different class assignments in another cell type, with ~6-13% falling in each of the MCO, -dREG, and, +dREG classes. Interestingly, +dREG and -dREG enhancers are commonly associated with promoter marks in a second cell type, likely reflecting an increase in the local cell-type-specific strength of Pol II initiation^19^. Together, these results support observations that promoters are more likely to be constitutive across cell types, whereas enhancers more frequently tend to be cell-type-specific and can transition between the classes of TRE.

### dREG Identifies H3K27ac-enriched “Active” Transcriptional Regulatory Elements

We investigated the distinctions among the four classes of TREs in more detail by comparing their genomic distributions with those of specific histone marks. In particular, we characterized the enrichments among the MCO, -dREG, +dREG, and Insulator classes of three histone marks—H3K27ac, H3K9ac, and H3K4me1—that have previously been reported to mark distinct classes of TRE. Whereas H3K27ac and H3K9ac denote ‘active’ regulatory elements^27,28^, H3K4me1 is a universal mark located at both active and so-called ‘primed’ enhancers^29^. We found that dREG TREs are strongly enriched for the ‘active’ H3K27ac and H3K9ac signals and, accordingly, that the vast majority of ENCODE peak calls for these marks are also identified by dREG. In contrast, the -dREG and MCO classes show little or no H3K27ac or H3K9ac signals (Fig. 4A,B; Supplementary Fig. 5). Moreover, the minority of +dREG TREs that are not associated with H3K27ac peak calls nevertheless display elevated H3K27ac ChIP-seq signals (Supplementary Fig. 6), suggesting that many simply fall below the detection threshold used in peak calling. This observation suggests that H3K27ac and +dREG point to the same class of functional element, in sharp contrast to previous work^30,31^. H3K4me1 is not only enriched at dREG TREs (both enhancers and promoters), but is also found at high levels in the -dREG and MCO classes (Fig. 4A-B). Thus, dREG identifies the same genomic regions as detected using ChIP-seq for H3K27ac and H3K9ac, and a subset of H3K4me1 peaks, suggesting that it can effectively distinguish between ‘active’ and ‘poised’ enhancer classes.

**Fig. 5:**
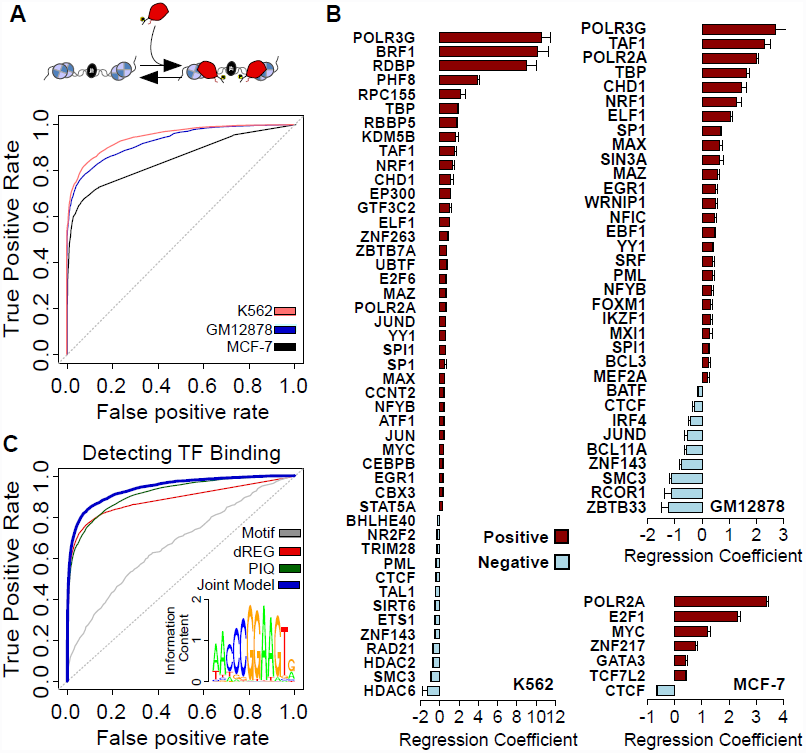
Sequence-specific transcription factors distinguish between DNase-I hypersensitive and transcribed regulatory TREs. (A) TFs are either associated with DNase-I hypersensitive peaks that are actively transcribed (+dREG) or open but non-transcribed (-dREG and DNase-I hypersensitive insulators), as indicated by the presence of Pol II (red rocket). ROC plots depict the accuracy with which these classes of regulatory TRE can be distinguished in three cell types based on the patterns of TF binding. (B) Logistic regression coefficients for each transcription factor correlated with transcription initiation (positive, red) or repression (negative, blue) following a 1,000 sample bootstrap. (C) Accuracy predicting TF binding (by ChIP-seq) using PIQ (green), dREG (red), the motif (gray), or a joint logistic regression model considering all three variables (blue).

**Fig. 6:**
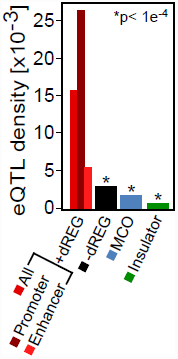
eQTL enrichments in the four classes of functional element. The density of eQTLs per site found in +dREG (further divided into promoters and enhancers using ChromHMM), -dREG, modified chromatin only (MCO), and Insulator classes. Labels appear in the same order as the associated bar, and colors shown below each label.

### Elements Marked Only by Chromatin are Occluded by a Central Nucleosome

The observation that TREs in the MCO class are generally not accessible to DNase-I cleavage suggests that access to binding sites in these TREs might be restricted by nucleosomes or higher-order forms of chromatin structure. To test this possibility, we used MNase-seq data to map the locations of nucleosomes surrounding all four classes of TREs in K562 cells. We found that TREs in the MCO class have a well-positioned nucleosome near their center (Fig. 4C), which likely occludes binding by transcriptional activators as well as cleavage by DNase-I. By contrast, -dREG enhancers typically contain an array of well-positioned nucleosomes in which the central nucleosome appears to have been displaced, whereas +dREG TREs, on average, contain a large nucleosome-free region surrounding the center and extending for ~1-2 kbp in both directions (although this pattern is more prominent at promoters than enhancers; see Supplementary Fig. 7). Thus, there is a fundamental distinction in nucleosome positioning between the MCO class and classes additionally characterized by DNase-I hypersensitivity and/or active transcription.

### Sequence-specific Transcription Factors and Co-factors Activate and Suppress eRNA Synthesis

The observation that nucleosomes occlude specific classes of TRE suggests that we might observe differences in the repertoires of sequence-specific TFs that bind TREs of each class. Therefore, we examined binding patterns for 91 TFs and co-factors for which high-resolution ChIP-seq data is available in K562 cells. We found that most TREs in the MCO class do not bind any sequence-specific TFs (Fig. 4D), and most -dREG and Insulator TREs bind small numbers of TFs (i.e., 1–10). dREG TREs, by contrast, display a striking enrichment for binding many TFs (39% bind more than 20 TFs). Within the +dREG set, promoters recruit the highest numbers of TFs, but transcribed enhancers are significantly more highly populated by TFs than TREs in the MCO and -dREG classes (e.g., 21% bind more than 20 TFs, compared with 5% for -dREG TREs). The same pattern was also observed in GM12878 and HeLa carcinoma cells, for which abundant ChIP-seq data is also available.

The transition between DNase-I accessible but non-transcribed elements (-dREG and some Insulators) and actively transcribed promoters or enhancers (+dREG) is likely to be specified by TFs that occupy active TREs. To identify TFs that might contribute to transcriptional activation at TREs we created a logistic regression model with the transcription status of each distal TRE as the response, and the presence or absence of ChIP-seq-assayed TF binding events within the TRE (in K562, GM12878, MCF-7, and HeLa cells) as the predictors. In this analysis, we augmented data from ENCODE with data from a recent ChIP-chip study profiling the binding of an additional 37 TFs in MCF-7 cells^32^. Our regression model predicts the transcription status of a holdout set of DHSs with remarkably high accuracy (Fig. 5A; AUC= 0.85-0.94). In comparison, a similar model that considers only the absolute level of DNase-seq signal intensity performs substantially worse (AUC= 0.78 in K562). These observations reveal that binding by particular TFs, more than simply the degree of chromatin accessibility, is responsible for the differential transcriptional outcomes observed in dREG TREs.

This regression analysis provides additional information about the relative importance of individual TFs in predicting whether or not a site is transcribed (Fig. 5B). A comparison of regression coefficients indicates, as expected, that components of the preinitiation complex, the histone acetyltransferase P300, and many sequence-specific activators (e.g., AP-1, PU1, CEBPB) are highly predictive of transcription initiation at TREs. By contrast, transcriptional co-repressors (e.g., HDACs and TRIM28), are associated with an absence of transcription. Notably, insulator-associated proteins (e.g., CTCF, RAD21, and SMC3) are also associated with an absence of transcription (Fig. 5B), consistent with the overlap observed between dREG sites and either CTCF peak calls (Fig. 4B) or raw signal (Supplementary Fig. 8). This finding strongly supports the distinction between Insulators and other classes of TRE. Unexpected cell-type-specific regulatory patterns also emerged from this analysis. For example, although FOS and JUN heterodimerize to form the AP-1 complex, they are associated with positive and negative regression coefficients, respectively, in GM12878. This curious pattern can be explained by the presence of BATF, another AP-1 member, which forms a heterodimer with JUN (but not FOS) and suppresses transcriptional activation by the AP-1 complex^33^. This finding strongly suggests that Jun-BATF heterodimers bind many AP-1 enhancers in GM12878 and suppress transcription activation. We conclude that transitions between –dREG and +dREG TRE classes are tightly regulated by a delicate balance of activation and active suppression mediated by specific combinations of TFs recruited to each site.

### Predicting Transcription Factor Binding using dREG

Having shown that TF binding is predictive of transcription initiation at TREs, we next addressed an inverse question: is transcription at TREs predictive of whether or not an individual TF is bound? Our interest here was in seeing whether dREG could be useful as a surrogate for, or complement to, DNase-seq data, which is widely used as an aid in the identification of cell-type-specific TF binding events^34–36^. As a proof of concept, we chose four transcriptional activators (NRF1, ELF1, SP1, and MAX) with a range of motif information contents^37^ and positive regression coefficients in the analysis described above. For all four TFs, we found that dREG scores alone perform nearly as well as the PIQ program^35^, which makes use of DNase-seq data in predicting ChIP-seq-supported binding events. In all cases dREG scores achieved a strikingly higher accuracy than matches of the DNA sequence to the motif alone. For example, for ELF1 (Fig. 5C), dREG produces a ROC score 3.4% lower than PIQ (AUC= 0.88 [dREG, red line] compared with 0.91 [PIQ, green line]) and both assays identify cell-type specific binding sites much better than motif matches alone (AUC= 0.66, gray line). Jointly modeling DNase-seq, dREG, and the motif match score improves classification accuracy 3.3-6.8% (Fig. 5C blue line; AUC= 0.94), substantially exceeding the PIQ score in this task. Thus, dREG appears to be a useful complement to DNase-seq based models of TF-DNA interaction for sequence-specific activators.

### Enrichment for eQTLs and GWAS hits in dREG Predictions

We hypothesized that dREG specifically marks the subset of open-chromatin sites that are actively regulating gene expression in the current cellular context. To explore this possibility, we compared the density of expression quantitative trait loci (eQTLs) identified in lymphoblastoid cell lines (LCLs)^38^ among +dREG, -dREG, and MCO TREs. We found that +dREG TREs in GM12878 LCLs contain 5.3–23.6-fold higher eQTL densities in LCLs than in other classes of TREs (Fig. 6), and account for 523 out of 818 of the eQTL SNPs that intersect with our set of TREs (~64%). Although this observation is partially explained by systematic biases in eQTLs density for gene promoters, if we focus on TREs associated with ‘enhancers’ only, we still observe a 1.9–8.2-fold enrichment in eQTL densities in +dREG TREs relative to the -dREG, MCO, and Insulator classes (p< 1e-4; Fisher’s Exact Test). This residual enrichment cannot be explained by differences in the distributions of the distance of these site classes relative to TSS annotations (Supplementary Fig. 9).

Next, we compared dREG predictions to a set of putatively functional single-nucleotide variants detected in a broad collection of genome wide association studies (GWAS)^39^. We found that dREG sites detected in relevant primary cell types are significantly enriched in GWAS-associated SNPs. For example, SNPs associated with autoimmune-related disorders are enriched in dREG sites in CD4+ T-cells and GM12878 LCLs (B-cells), including SNPs for celiac disease (11.3 and 9.7-fold, respectively), rheumatoid arthritis (8.4 and 11.9), and type-1 diabetes (6.2 and 7.1). Similarly, SNPs associated with asthma are 9.7-fold enriched in B-cells (but were not enriched in T-cells). Thus, both eQTLs and GWAS associations are significantly enriched in the transcribed regulatory elements identified by dREG, suggesting that these TREs are more likely to be actively regulating gene expression than other classes of TRE.

### Discussion

In this article, we have introduced a new high-throughput prediction method, called dREG, for detecting active transcriptional regulatory elements (TREs) from patterns of RNA polymerases evident from standard GRO-seq or PRO-seq data. We show that dREG can achieve both high sensitivity (>90%) and high specificity (genome-wide FDR < 10%) for active TREs identified by GRO-cap and CAGE^19^. Our approach allows the identification of many types of functional elements that play a vital role in gene activation, including TREs, TF binding, gene expression, pausing levels, and elongation rates, to be measured using a single PRO-seq experiment. This efficiency is vital in a number of exciting applications, for example in cancer genomics and personalized medicine, in which the application of genomics technologies are currently limited by sample quantities and the high cost of collecting data in large numbers of subjects.

By comparing dREG sites to other functional genomic assays, we demonstrate the existence of at least four major classes of TREs in human cells. These classes correspond to closed chromatin marked by chromatin modifications such as H3K4me1 (MCO), DNase-I accessible DNA without a dREG signal (-dREG), insulator factor binding (CTCF), and divergent transcription detected by dREG (+dREG). Several lines of evidence, including enrichments for eQTLs, transcriptional activators, and histone acetylation strongly support our proposal that dREG identifies the genomic sites that play a direct and active role in gene regulation.

We discovered three independent classes of regulatory elements that are untranscribed (-dREG, MCO, and Insulator). These TREs both appear to be inactive by a variety of metrics, including a paucity of eQTLs (Fig. 6A), depletion of transcription factor binding (Fig. 4D), and the absence of histone acetylation (Fig. 4A,B). We think that insulators are a distinct functional class, as they are depleted for the functional marks examined here, yet appear to constitute a complete functional element^40^. These observations suggest that CTCF likely has a distinct functional role in maintaining the three-dimensional architecture of DNA^41^, but can indirectly affect gene expression under certain circumstances^40^.

The second class of inactive sites (MCO) is characterized by a subset of the chromatin modifications associated with regulatory regions (e.g., H3K4me1). These TREs are generally devoid of transcription factor binding (Fig. 4D), and the majority are not DNase-I hypersensitive (Fig. 3A). Interestingly, on average, a nucleosome overlaps the center of these sites (Fig. 4C), which apparently competes with transcriptional activators for DNA binding. These sites appear similar in many respects to DNase-seq-depleted, H3K4me1-enriched peaks that nevertheless appear to drive transcriptional activation *ex vivo*, as determined using STARR-seq, a high-throughput reporter gene assay^6^. Taken together, these observations are consistent with a model in which the DNA sequence in these regions can drive transcriptional activation in a permissive chromatin context, but the binding of certain TF activators is prevented by steric competition with chromatin at native sites.

The third class of inactive site (-dREG) is characterized by high-confidence DNase-I accessibility and similar patterns of nucleosome positioning to dREG enhancer elements, but an absence of transcription initiation. These sites are, on average, more likely to be bound by HDACs and repressive transcription factors than +dREG sites (Fig. 5B), suggesting that transcription is actively suppressed. These observations suggest that the local chromatin context is configured in a manner that might permit transcription, but transcriptional activation is being actively suppressed by the cell. Thus, we identify at least two separate mechanisms by which regulatory regions may be inactivated, including the presence of a central nucleosome that occludes activator binding (epigenetic), and DNA sequence-dependent suppression at open chromatin (genetic). These observations demonstrate that numerous genetic and epigenetic conditions must be met in order to construct an active regulatory element.

The specific mechanisms by which multiple TFs cooperatively regulate gene expression remain incompletely understood at a genome-wide level. Our observation that at least two mechanisms are responsible for silencing TREs suggests that one requirement for multiple TFs is to overcome the multiple, independent mechanisms by which regulatory elements are inactivated. Several lines of evidence support this idea. First, we observe that transcribed TREs tend to be associated with larger numbers of TFs than inactive TREs (Fig. 4D). Second, the presence of certain TFs removes functional marks from TREs that correlate with patterns of inactivity that we observe here (e.g., ‘pioneer’ TFs such as FOXA1 and GATA3 prepare the chromatin environment for TF binding^35,42,43^, in some cases by evicting a competing nucleosome^44^). Third, inactive TREs can undergo rapid, stimulus-dependent transitions from either –dREG or MCO to +dREG^14,44^. These observations suggest that individual TFs are able to overcome some (but often not all) of the suppressive local effects at TREs, and indicate that multiple TFs work together to construct an active TRE.

Transitions between three classes, namely MCO, -dREG, and +dREG, following stimulation or across different cell types indicates that they are fundamentally related to one other. We hypothesize that these classes of functional element represent intermediate states in either the assembly or disassembly of TREs. Notably, although MCO and -dREG are depleted for cell-type specific eQTLs, patterns of evolutionary conservation are relatively consistent across the different classes and universally higher than expected under neutral drift (Supplementary Fig. 10). This suggests that all classes are enriched for functional elements that are active in at least one cell type. Thus, our data suggests that MCO, -dREG, and +dREG classes of TRE represent intermediate states in either the assembly or disassembly of active regulatory elements. We expect that future studies will address the specific functional mechanisms that underlie these transitions, and will further elucidate the relationship between these important classes of functional element.

## Methods

### Training the Support Vector Regression Model

#### Overview

We consider transcription start site detection using GRO-seq and PRO-seq (hereafter we refer only to GRO-seq, but the same methods apply to both sources of data) data as a regression problem. Our goal is to separate regions of high GRO-seq signal intensity into a class in which RNA polymerase originates by initiation and rapidly transitions to elongation (positive set, comprised of transcription start sites), and a class that polymerase elongates through (negative set, largely comprised of gene bodies). This classification problem was addressed using a standard epsilon-support vector regression (SVR), as described in the following sections.

#### GRO-seq Signal Intensity Requirements

We removed from consideration any loci with very low signal levels, implicitly assigning these positions to the negative set. We retained sites meeting either of the following two signal intensity thresholds: one or more reads on both the plus and minus strand within a window of 1 kbp, or three or more reads within a window of 100 bp on either the plus or minus strand. At these cutoff thresholds, 93% of K562 GRO-cap peaks contain at least one informative site in a PRO-seq library depth of ~40M reads. The remaining sites were segmented into non-overlapping 50 bp intervals to improve the speed of processing on large datasets.

#### GRO-seq Feature Vector

GRO-seq read counts are summarized in our model as a multi-scale feature vector, as illustrated by the barchart in Fig. 2A. We count GRO-seq reads that map in non-overlapping windows on either side of a central base that meets the signal intensity requirements (as described above). Our approach represents the genome at multiple scales (window sizes). For each scale, we count reads in the specified number of non-overlapping windows both upstream and downstream of the central base. Each scale can represent redundant information in the GRO-seq read counts. The final feature vector is constructed by concatenating the vectors representing read counts at each scale and strand. The specific parameters of the scales and number of windows at each scale were optimized using cross-validation (as described below, and depicted in Supplementary Table 2).

#### Data Standardization

GRO-seq data was standardized using the logistic function, *F*(*t*), with parameters α and β, as follows:

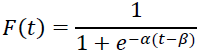

where *t* denotes the read counts in each window. We find it convenient to define the `tuning’ parameters *α* and *β* in terms of a transformed pair of parameters, *x* and *y*, such that *x* represents the fractional portion of the maximum read count depth at which the logistic function reaches 1 and *y* represents the value of the logistic function at read counts of 0. The relationship of (*α*, *β*) to (*x*, *y*) is given by the following equations:

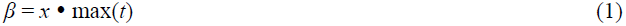

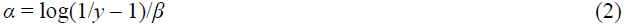

where max(*t*) denotes the maximum read depth, as computed separately for each window size and strand in the feature vector. In practice, we find it convenient to fix the value of *y* at 0.01 and use *x* for tuning. We tried values of *x* between 1% and 100% of the maximum read depth, and found that the optimal AUC was achieved at *x* = 0.05 (Supplementary Table 1). Using this function in its optimized form will generally assign each position values near 0 or 1, and consequently most of the signal for dREG is dependent on where reads are located, rather than on the relative read depths.

We also evaluated alternative standardization approaches, including simply dividing the reads in each feature vector by their maximum value, but these approaches did not perform as well as the logistic function.

#### Training the dREG Support Vector Regression model

We fit an epsilon-support vector regression model using the e1071 R package^45^, which is based on the libsvm SVM implementation^46^. When training dREG, we assigned a label of 1 to sites intersecting GRO-cap transcription start sites^19^, and excluded from the training set any sites intersecting a functional mark indicative of a regulatory element but not a GRO-cap peak (including ChromHMM enhancers or promoters). All other positions in the genome meeting the GRO-seq signal requirements (described above) were assigned a score of 0. The final SVR was trained on a matched set of 100,000 loci using PRO-seq data in K562 cells. Sites in the positive set (i.e., GRO-cap peaks) were chosen at random. When selecting the set of negative (i.e., non-transcription start site) examples, we chose 25% of sites to enrich for positions that were commonly associated with false positives during preliminary testing. These include 15% of the negative set that were selected to be within 1-5 kbp of the positive regions (to improve the resolution of dREG), and 10% in regions where the 3' ends of annotated genes on opposite strands converged (to eliminate a common source of false positives). The remaining 75% of negative sites were selected at random from the set of positions across the genome meeting the GRO-seq signal requirements (described above).

#### Optimizing Tuning Parameters

Tuning parameters were optimized on a balanced set of 50,000 loci, and performance was evaluated on a holdout set of 2,000. Parameters were chosen to maximize the area under the receiver operating characteristic curve (AUC). We first selected parameters of the data transformation that maximized the AUC using a fixed feature vector (20 windows, each 10, 50, and 500 bp in size). Subsequently we fixed the optimal data standardization (see *Data Standardization* section, above) and selected the feature vector, including the number and size of windows, which maximized the AUC. False positives are defined as sites which do not overlap GRO-cap, DHSs, or ChromHMM (promoters, enhancers, or insulators). True positives are sites that overlap GRO-cap HMM predictions^19^. False negatives are sites that are identified by GRO-cap, but are not identified by dREG. True negatives are sites that are not identified by dREG, or any of the other assays. Various tuning parameter settings are summarized in Supplementary Tables 1-2.

#### Running dREG and Post Processing

We ran dREG on GRO-seq or PRO-seq data in eight cell types. We used the SVR model trained in K562 cells to compute the predicted score at each position meeting the GRO-seq signal intensity thresholds. To call dREG ‘peaks’ we threshold this score at 0.8, which we found returned a 10% false discovery rate (FDR) in two datasets for which extensive data was available (K562 and GM12878). In cell types with lower read counts, this score is likely to be conservative, resulting in both a lower FDR, and lower sensitivity (see Supplementary Fig. 1). Regions meeting the dREG signal requirement within 500bp of one another were merged to prevent spurious detection of dREG sites adjacent to bona-fide TREs, or independent detection of the same promoter or enhancer elements.

#### dREG Sensitivity to Sequencing Depth and Library Quality

To evaluate the sensitivity of dREG to sequencing depth we subsampled the K562 data by removing reads at random from the bed files representing mapped reads. We ran the dREG algorithm as described, either with or without re-training the model on the reduced read depth (both are plotted in Supplementary Fig. 1). Artificial low quality datasets were created by randomly dropping out some locations and redistributing their reads to neighboring sites in a 50 kbp (non-overlapping) window. In each window, locations were retained with probability proportional to the original read density at that site. This procedure was designed to re-create the profile observed in low-quality data, in which large numbers of reads tend to align on a small number of positions, creating the appearance of ‘spikes’ when viewed on the genome browser. The asymptotic unique reads metric used to evaluate data quality was defined as the number of unique genomic coordinates in a GRO-seq library as the number of mapped reads approaches infinity. This value was estimated by subsampling the read depth of the data and recording the number of unique locations covered. By observing the critical point in this function (where a numerical approximation to the derivative was close to zero, defined as within 1% of the final value) we could infer the point at which the library had 'saturated' its coverage of unique locations. This number of unique locations was then taken to be the asymptotic unique reads. When necessary, this value was interpolated under the assumption that the slope of the read depth does not change.

### GRO-seq and PRO-seq Library Prep

#### Extraction of Primary CD4+ T-cells from Blood Samples

Blood samples (80-100mL) from three human individuals were collected at Gannett Health Services in compliance with Cornell IRB guidelines. Mononuclear cells were isolated using density gradient centrifugation, and CD4+ cells were extracted using CD4 microbeads from Miltenyi Biotech (130-045-101), following the manufacturer’s instructions. Primary CD4+ T-Cells were kept in culture (RPMI-1640, supplemented with 10% FBS) for 1-3 hours to recover homeostasis.

#### Cell Culture Conditions and PRO-seq Library Preparation

Both primary and Jurkat CD4+ T-cells were maintained in RPMI-1640 media supplemented with 10% FBS, and treated for 30 minutes with low amounts of DMSO and ethanol (as they are controls for a separate experiment, manuscript in preparation). To isolate nuclei, cells were resuspended in 1mL lysis buffer (10mM Tris-Cl pH 8, 300mM sucrose, 10mM NaCl, 2mM MgAc2, 3mM CaCl2, and 0.1% NP-40). Nuclei were washed in 10mL of wash buffer (10mM Tris-Cl pH 8, 300mM sucrose, 10mM NaCl, and 2mM MgAc2) to dilute free NTPs. Nuclei were washed in 1mL, and subsequently resuspended in 50uL, of storage buffer (50mL Tris-Cl pH 8.3, 40% glycerol, 5mM MgCl2, and 0.1mM EDTA), snap frozen in liquid nitrogen, and kept for up to 6 months before preforming PRO-seq. HeLa cells were maintained in DMEM media supplemented with 10% FBS and 1x pen/strep (Gibco). Cells were harvested by rinsing the tissue culture plate several times in 1x PBS followed by scraping in 10ml of 1x PBS. Cells were pelleted by centrifugation and nuclei were isolated as described above. K562 cells were maintained in culture and nuclei were isolated exactly as previously described^19^. For all cell types, PRO-seq or GRO-seq was performed as exactly described^10,16^, and sequenced using an Illumina Hi-Seq 2000 at the Cornell University Biotechnology Resource Center.

### Comparison to ChromHMM and DNase-I data

We compared dREG TREs to ENCODE DNase-I and ChromHMM data. For ChromHMM data, we selected the set of sites annotated as promoter, enhancer, or insulator using data from GM12878, K562^24^, HeLa^47^, or CD4+ T-cells^48^. We collected ENCODE DNase-I peak calls from the UW or Duke DNase-I-seq protocol^2^, and selected peaks identified using both experimental assays. To compare across different experimental assays, we merged sites identified by ChromHMM, DNase-I-seq, and dREG, and labeled each merged site based on the experimental assays which identified it. TREs were subsequently divided into four independent, non-overlapping classes based on the set of experimental peak calls that they intersected. Site classes were defined as those sites that intersect: (1) dREG, DNase-I-seq, and ChromHMM (+dREG), (2) ChromHMM insulators but not dREG (Insulator), (3) DNase-I-seq and ChromHMM, but not dREG (-dREG), and (4) ChromHMM, but not DNase-I-seq or dREG (Modified Chromatin Only; or MCO). All operations in these analyses were performed using the bedops^49^, bedtools^50^, or bigWig software packages.

### Logistic Regression Classifier of DNase-I peaks with and without dREG

We used a logistic regression classifier to evaluate the how accurately transcription factors (TFs) could be used to distinguish between DNase-I peaks with and without the presence of dREG. We collected the set of all high confidence DNase-I peaks, consisting of the intersection between the UW and Duke assays. To improve our confidence about the transcription status of each DNase-I peak, we required that dREG-positive sites contain dREG scores greater than 0.8, and dREG-negative sites have a score was less than 0.3.

We modeled the presence or absence of dREG at a particular DNase-I peak as the response in a logistic regression. Co-variants consist of the presence or absence of each TF assayed in the cell type of interest. To determine the presence or absence of each TF, we collected uniform peak calls for all ChIP-seq data from the ENCODE project. For MCF-7 cells, ENCODE data was supplemented with a set of 37 TFs for which ChIP-chip data was available^32^. TFs having multiple ChIP biological replicates were associated with each peak if any of the replicates was enriched at that peak. The significance of the direction of effect for each TF on the presence of a dREG signal was determined using a 1,000-sample bootstrap, in which we chose one TF at random to omit from the regression analysis during each iteration. Fig. 5B plots the set of all TFs for each cell type whose direction of effect is consistent across each of the bootstrap iterations.

### Identification of TF Binding using dREG, DNase-I and a joint model

We identified all occurrences of motifs associated with four transcription factors (NRF1, ELF1, MAX, and SP1) in hg19 using a permissive motif-match log-likelihood ratio threshold of 5 for the motif relative to the background set. Each position was classified as ‘bound’ or ‘unbound’ to the TF of interest using on ENCODE ChIP-seq peak calls in the appropriate cell type. ROC plots profiling the accuracy of binding detection were collected by varying the max dREG score in a 200 bp window (treating unscored sites as a score of 0), DNase-I read counts in a 200 bp window around each putative motif matching the canonical PWM, or more stringent matches to the canonical TF motif. PIQ was run using the instructions provided by the authors. To evaluate the accuracy of PIQ we varied the threshold of the predicted positive-predictive value output by the PIQ program at each site. We also evaluated a joint model which used data from dREG, DNase-I, the motif, and the absolute amount of Pol II mapping to the forward and reverse strand (within 200 bp) using each data source as a covariate in a logistic regression, modelling the presence of a ChIP-seq peak at each motif match as the response variable.

## Acknowledgements

We think Iris Jonkers and Noah Dukler for comments and helpful discussions on an early manuscript draft, and Brad Gulko for critical discussions about support vector machines. This work was made possible by generous seed grants from the Cornell University Center for Vertebrate Genomics (CVG), Center for Comparative and Population Genetics (3CPG), an NHGRI grant (5R01HG007070-02) to ACS and JTL, and NIH R01 (DK058110) to WLK.

## Author Contributions

CGD designed the dREG tool. CGD, ALM, and SLH designed and implemented the software. CGD, SLH, ALM, LJC, JTL, and AS analyzed the data and interpreted the results. LJC, CTW, CGD, HWL, JTL, WLK, and VGC contributed previously unpublished data, helped troubleshoot experiments, and/or preformed library preps. CGD, AS, JTL, LJC, SLH, and ALM wrote the manuscript.

## Figure Legends

**Supplementary Fig. 1:**
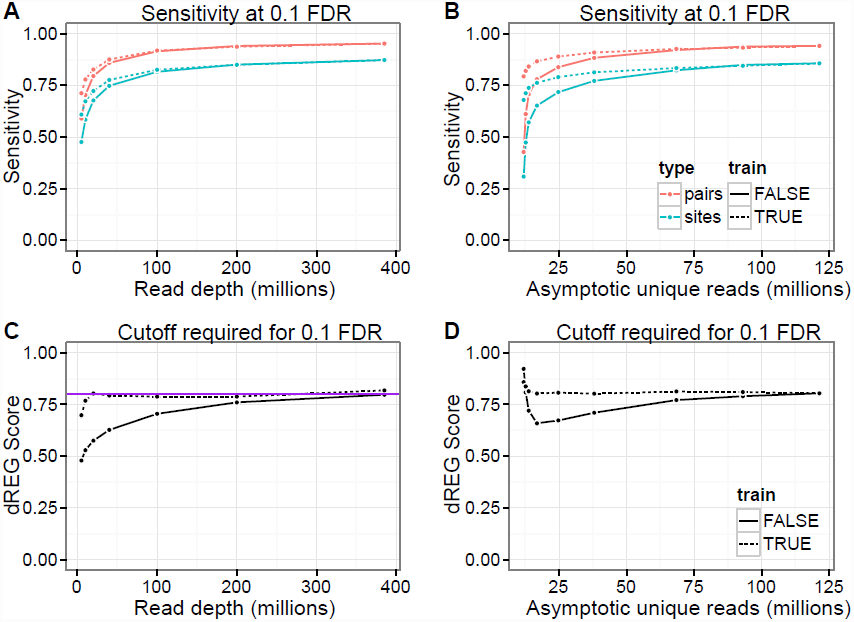
Sensitivity to read depth and library quality. (A) dREG sensitivity at a 10% false discovery rate at the indicated read depth or asymptotic library complexity. Dotted lines indicate a model that has been trained specifically on the indicated library. Solid lines indicate the model trained on the native K562 PRO-seq libraries. Pink and cyan denote GRO-cap sites and pairs, respectively. (C-D) SVR threshold required to achieve a 10% FDR for SVR models that have (dotted) or have not (solid) been trained specifically to the indicated parameters.

**Supplementary Fig. 2: dREG TREs are associated with chromatin marks characteristic of both promoters and enhancers.**
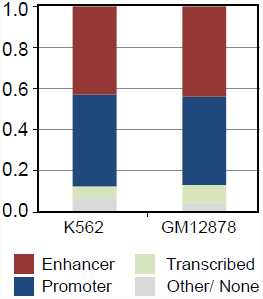
The majority (>90%) of dREG TREs intersect post-translational histone modifications previously associated either promoters or enhancers, and interpreted by ChromHMM.

**Supplementary Fig. 3:**
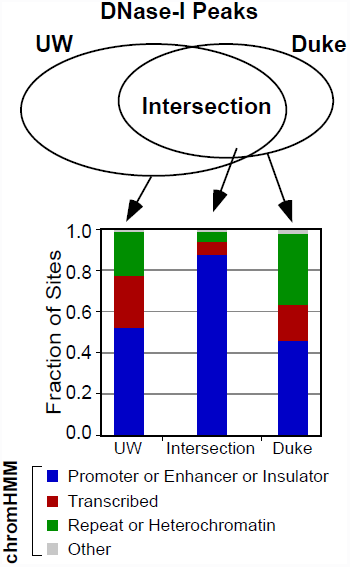
Chromatin marks associated with three classes of DNase-I hypersensitive sites. DNase-I hypersensitive sites identified by either the UW and Duke assays alone, or their intersection, are associated with the indicated fraction of regulatory marks (blue), transcribed regions (red), or repeat/heterochromatin (green), as annotated by ChromHMM.

**Supplementary Fig. 4:**
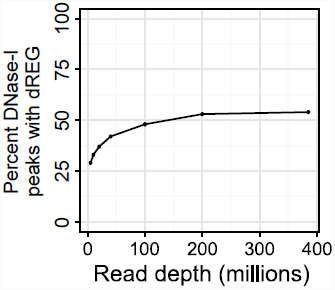
High-confidence DNase-I Peaks covered by dREG. Fraction of DNase-I peaks (excluding CTCF-bound insulators) that intersect a dREG site (Y-axis) as a function of the PRO-seq read depth (X-axis) in K562 cells.

**Supplementary Fig. 5:**
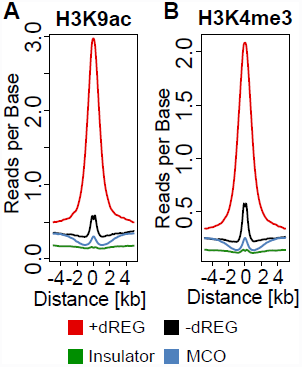
Association of TREs in each class to independent functional marks. Comparison of read-densities for H3K9ac (A) and H3K4me3 (B) in each class of functional element.

**Supplementary Fig. 6:**
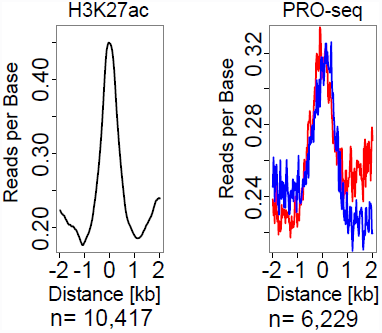
Enrichment of H3K27ac and PRO-seq signal intensity. Enrichment of H3K27ac at dREG TREs that lack an H3K27ac peak call (left); and PRO-seq signal on the plus (red) and minus (blue) strand at H3K27ac peak calls without a dREG TRE prediction.

**Supplementary Fig. 7:**
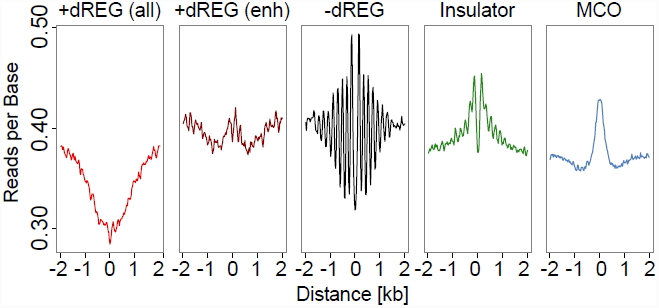
MNase-seq signal near distinct classes of TRE. Plots show the ENCODE MNase-seq signal centered on the indicated class of TRE.

**Supplementary Fig. 8:**
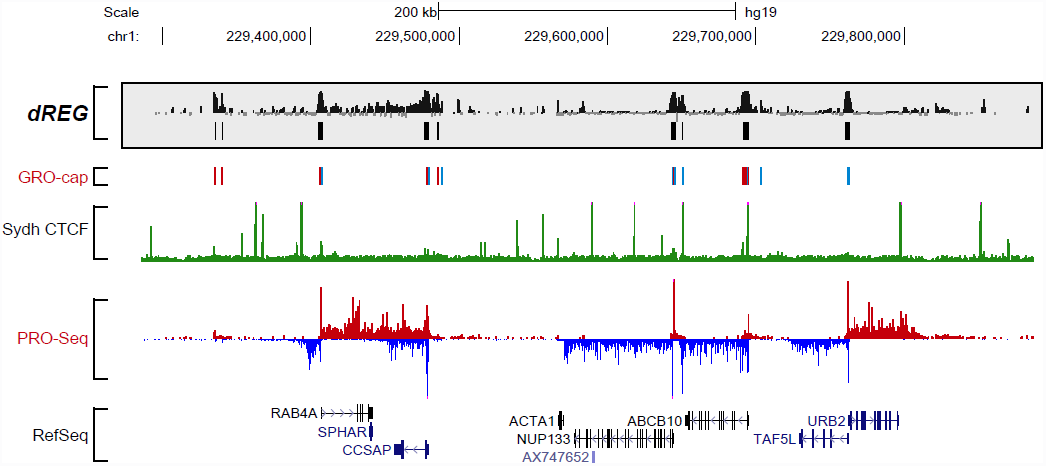
Browser shot of CTCF ChIP-seq and PRO-seq signal. UCSC genome browser signal compares dREG, GRO-cap, CTCF, and PRO-seq in the indicated region of chromosome 1 in K562 cells.

**Supplementary Fig. 9:**
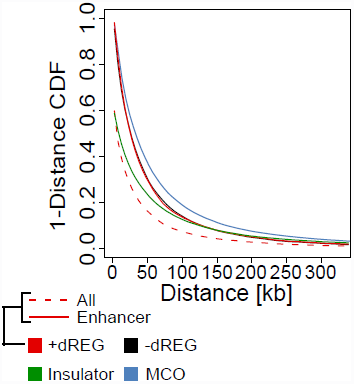
Distance of each class to the nearest RefSeq annotated transcription start site. Each point shows the fraction of TREs in the indicated class with a distance to the nearest RefSeq annotated transcription start site greater than the value indicated on the X-axis (i.e., 1-cumulative density function). In this plot, separate lines show the distribution for the set of all +dREG TREs (red, dotted) and for the subset which intersects chromatin marks indicative of enhancers (red, solid).

**Supplementary Fig. 10:**
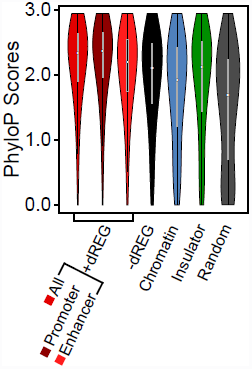
PhyloP scores among the placental mammals in each class of TRE. Violin plots denote the distribution of the maximum PhyloP score within each occurrence of the indicated class of TRE in GM12878 cells.

## Tables

**Supplementary Table 1:** Optimization of regularization of PRO-seq and GRO-seq data. We optimized the regularization/data normalization procedures on a simple, fixed feature vector. The accuracy (measured by AUC) of each standardization procedure are shown in the indicated row of the table.

| Transform | $x$ | AUC |
| --- | --- | --- |
| Logistic | 0.50 | 0.9195 |
| Logistic | 0.25 | 0.9309 |
| Logistic | 0.20 | 0.9345 |
| Logistic | 0.15 | 0.9388 |
| Logistic | 0.10 | 0.9438 |
| Logistic | 0.05 | 0.9494 |
| Logistic | 0.01 | 0.9485 |

**Supplementary Table 2:** Optimizing the multi-scale feature vector. We optimized the multi-scale feature vector on a training set of 50,000 matched transcription start sites and control regions. Optimization was performed after fixing the data regularization settings at an optimal value on a simple feature vector.

| Window Sizes | Number of Windows | AUC |
| --- | --- | --- |
| 10, 50, 500 | 20, 20, 20 | 0.958557 |
| 10, 50, 500, 5000 | 20, 20, 20, 20 | 0.962113 |
| 25, 50, 500, 5000 | 20, 20, 20, 20 | 0.961917 |
| 10, 25, 50, 500, 5000 | 20, 20, 20, 20, 20 | 0.962759 |
| 10, 50, 500, 5000 | 30, 20, 20, 20 | 0.960941 |
| 10, 50, 500, 5000 | 20, 30, 20, 20 | 0.962276 |
| 10, 50, 500, 5000 | 20, 20, 30, 20 | 0.961133 |
| 10, 50, 500, 5000 | 20, 20, 20, 30 | 0.962235 |
| 10, 50, 500, 5000 | 10, 20, 20, 20 | 0.964225 |
| 10, 50, 500, 5000 | 20, 10, 20, 20 | 0.961931 |
| 10, 50, 500, 5000 | 20, 20, 10, 20 | 0.962623 |
| 10, 50, 500, 5000 | 20, 20, 20, 10 | 0.961765 |
| 50, 500, 5000 | 30, 20, 20 | 0.959634 |
| 10, 50, 500, 5000 | 10, 30, 20, 20 | 0.963887 |
| 10, 50, 500, 5000 | 15, 30, 20, 20 | 0.963370 |
| 10, 25, 50, 500, 5000 | 10, 10, 30, 20, 20 | 0.964326 |

**Supplementary Table 3:**
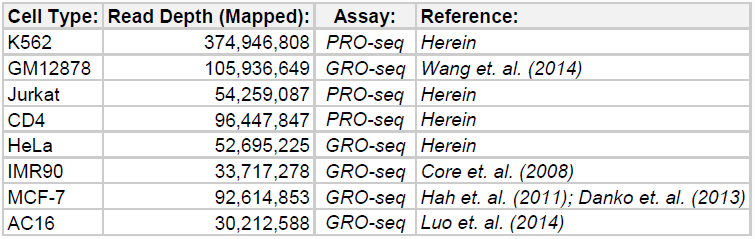
Libraries analyzed across eight cell types. Table columns show the cell type, number of mapped reads, assay type, and reference for the eight cell-types analyzed.

| Cell Type: | Read Depth (Mapped): | Assay: | Reference: |
| --- | --- | --- | --- |
| K562 | 374,946,808 | PRO-seq | Herein |
| GM12878 | 105,936,649 | GRO-seq | Wang et. al. (2014) |
| Jurkat | 54,259,087 | PRO-seq | Herein |
| CD4 | 96,447,847 | PRO-seq | Herein |
| HeLa | 52,695,225 | GRO-seq | Herein |
| IMR90 | 33,717,278 | GRO-seq | Core et. al. (2008) |
| MCF-7 | 92,614,853 | GRO-seq | Hah et. al. (2011); Danko et. al. (2013) |
| AC16 | 30,212,588 | GRO-seq | Luo et. al. (2014) |

